# Multiscale Molecular Modelling of Chromatin with MultiMM: From Nucleosomes to the Whole Genome

**DOI:** 10.1101/2024.07.26.605260

**Authors:** Sevastianos Korsak, Krzysztof Banecki, Dariusz Plewczynski

## Abstract

**Motivation:** We present a user-friendly 3D chromatin simulation model based on OpenMM, addressing the challenges posed by existing models with use-specific implementations. Our approach employs a multi-scale energy minimization strategy, capturing chromatin’s hierarchical structure. Initiating with a Hilbert curve-based structure, users can input files specifying nucleosome positioning, loops, compartments, or subcompartments.

**Results:** The model utilizes an energy minimization approach with a large choice of numerical integrators, providing the entire genome’s structure within minutes. Output files include the generated structures for each chromosome, offering a versatile and accessible tool for chromatin simulation in bioinformatics studies. Furthermore, MultiMM is capable of producing nucleosome-resolution structures by making simplistic geometric assumptions about the structure and the density of nucleosomes on the DNA.

**Code availability:** Open-source software and the manual are freely available on https://github.com/SFGLab/MultiMM.

## 1. Introduction

Chromatin, exhibiting a multi-scale effect (1), comprises nucleosomes as its fundamental units (1 *pb* − 10 *kb*), consisting of 8 histones that interact with DNA through electromagnetic forces (2). At a higher scale (50 *kb* − 200 *kb*) (3), loops form, driven by Smc complexes with ring-like topology acting as loop extruders and CTCF proteins serving as extrusion barriers (4). Differential densities of loops and nucleosomes induce a phase separation, leading to the compartmentalization of chromatin into two distinct compartments (3 *Mb*), a phenomenon extendable to four or five subcompartments (5). Further complexity arises from the interaction of chromosomes (100 *Mb* − 3000 *Mb*), each predominantly interacting with itself, within the genome (6). Notably, the nuclear lamina introduces an additional layer of intricacy, as specific chromatin regions interact with it (7, 8). Consequently, the hierarchical structure of chromatin implies different biophysical laws applicable at particular scales.

The resolution-specific impact of experimental data affects the complexity of chromatin analysis. Experiments like MNase-Seq or ATAC-Seq can determine the positioning of nucleosomes, and they can make a distinction between open and closed chromatin (9). Loop determination relies on 3C-type experiments like Hi-C, ChIA-PET, or Hi-ChIP, which give us information about the genome-wide interactions. At broader scales, compartmentalization patterns are revealed by analyzing the sign of the first eigenvector in Hi-C data (5), while more advanced models like Calder are able to classify chromosomes into subcompartments (10). Chromosomal territories naturally emerge from 3C-type experiments, reflecting a limited amount of intra-chromosomal interactions.

Modeling various scales of chromatin structure poses a formidable challenge due to the enigmatic nature of biophysical forces within chromatin. Nucleosomes, with a relatively higher degree of understanding, have been extensively studied through simulations capturing histone interactions with DNA mediated by electromagnetic forces (2). However, integrating these models for large chromatin regions is computationally impractical, and this is the reason why geometrical approaches for the simulation of nucleosomes have been developed (11). Stochastic models at the loop scale incorporate a rebinding mixing probability for Smc complexes diffusing in one dimension, hindered by CTCF barriers (4). In genome-wide models, fixed harmonic bond forces (12) are often assumed to circumvent the computational complexities of Monte-Carlo simulations, with equilibrium lengths and spring coefficients fine-tuned based on loop strength or PET-count from Hi-C, Hi-ChIP, or ChIA-PET data. Compartmentalization forces are frequently represented using a block-copolymer model (13), simulating long-range attractive interactions between compartments sharing the same compartment. Exploring interactions with the nuclear lamina (7, 14), such as the attraction of the B-compartment near the lamina or the increased density of small chromosomes near the nucleus, provides insights into more realistic structures.

In this study, we introduce a streamlined and accessible model, MultiMM (Multiscale Molecular Modelling), capable of encompassing chromatin’s various scales. Our model efficiently reconstructs the 3D chromatin structure in minutes, requiring only minimal input data. Additionally, we achieve nucleosome resolution by employing geometric modeling, thus circumventing the need for time-intensive molecular dynamics simulations at this scale.

**Figure 1:**
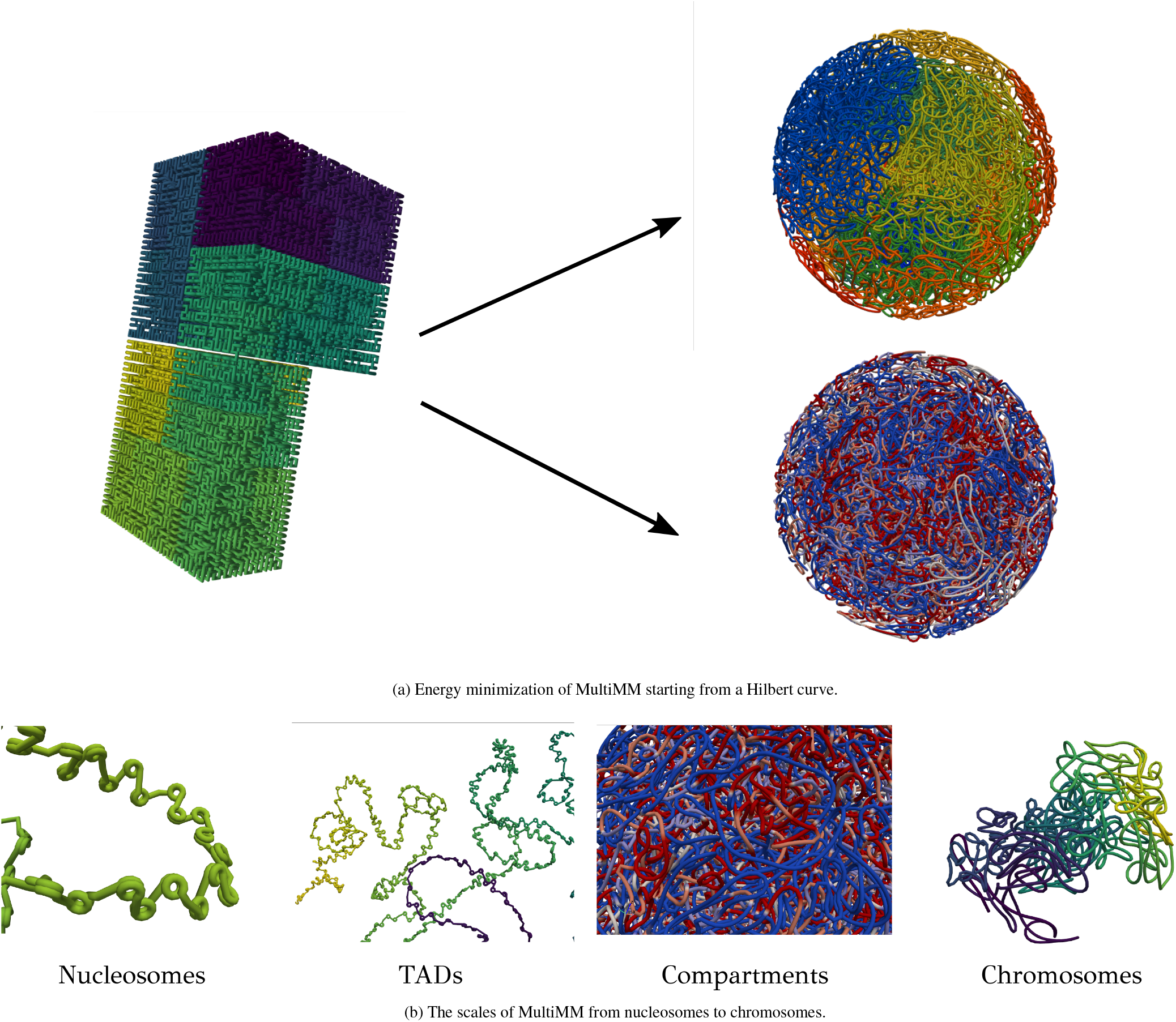
Simulation example with 50000 beads for GM12878. (a) We see how the Hilbert curve minimizes to genome-wide structures of chromatin, with chromosomes (up) and and compartments (down) for coloring. (b) The different scales of MultiMM. In the given figure, we can see nucleosomes, TADs, which represent groups of loops, compartments, which are consisted from groups of TADs, and the final structure of a chromosome.

## 2. Materials and Methods

### 2.1 Input parameters and data

The models are generated using a minimal dataset, which requires a file containing loops and the specification of coordinates and chromosomes for the region of interest. The simulation granularity is adjustable, allowing users to determine the number of beads utilized in the simulation. Optionally, users may input compartment or subcompartment files to model block-copolymer attractions. Nucleosomes are depicted as static helical structures, integrated into the final structure with densities corresponding to their ATAC-Seq or MNase-Seq signals. While MultiMM assumes default biophysical parameters for chromatin, users can modify these parameters via a configuration file, including the option to enable or disable forces. This flexibility makes MultiMM a user-friendly application capable of testing multiple hypotheses for chromatin folding. The simulation parameters and input data can be adjusted by editing a configuration file. Our model has been tested on various machines, demonstrating compatibility with laptops, workstations, and HPC clusters, and it can be parallelized in CPU or GPU according to the design of OpenMM (15).

### 2.2 Implementation of the model

In its initial steps, MultiMM adapts the provided loop data to match the simulation’s granularity, downgrading the data accordingly. It consolidates loop strengths by summing those associated with the same loop after downgrading and retains only statistically significant ones, applying a threshold value (so as to have minimum genomic distance 3 in simulation bead units). Loop strengths are then transformed to equilibrium distances by assuming an inverse power law relationship. Compartmentalization data are imported from Calder (10), and they are converted into a discrete value vector, where positive values represent A compartment and negative B compartment. In the case of nucleosomes, we assume that their density is inversely proportional to the ATAC-Seq signal, where a user-determined maximum amount of nucleosomes is assigned to each bead.

MultiMM constructs an initial structure based on a Hilbert curve after importing user-provided input data. The self-avoiding nature and inherent compaction of this initial structure provide significant advantages, notably expediting the energy minimization process. Each chromosome is represented by a distinct polymer chain, with interactions between different chromosomal chains being permitted. The initial positions of chromosomes on the Hilbert curve can be randomized by shuffling their starting and ending points within the initial structure. Subsequently, a multi-scale molecular force-field is employed, encompassing strong harmonic bond and angle forces between adjacent beads, along with harmonic spring forces of variable strength to model the imported long-range loops. For compartment modeling, a long-range block-copolymer potential with discrete energy levels is applied, where stronger forces denote robust attractive interactions between B-compartments. The number of energy levels corresponds to the count of distinct (sub)compartments. Intra-chromosomal interactions are also modeled as long-range harmonic bond forces, with the user responsible for providing relevant interactions to influence the resultant structure. The genome is folded within a spherical container with radii *R*_2_ = λ*R*_1_, λ > 1, where the smaller radius represents a boundary condition for the nucleolus. Addressing lamina interactions is challenging due to limited access to publicly available Lamin-Seq data. Consequently, we model these interactions by assuming that the B compartment is primarily attracted to the lamina, where *r* = *R*_1_ or *r* = *R*_2_. To represent this, we utilize a (sub)compartment-specific trigonometric potential with minima near the torus walls for the B-compartment and in the intermediate region for the A-compartment. Additionally, acknowledging that smaller chromosomes tend to be closer to the nucleolus than larger ones, we incorporate an attractive potential proportional to the chromosomal size and inversely proportional to the distance from the center. By minimizing the energy of this force-field by using an OpenMM integrator of preference, we can conclude to genome-wide structures.

For the implementation of the model, the python front-end of OpenMM was used (15). The energy minimization can be accelerated across multiple CPUs, or with platforms CUDA and OpenCL. Appropriate force-field parameters are selected by default, though the users may change them according their special preferences.

### 2.3 Resulting files

Following the execution of MultiMM, a file containing the minimized genome-wide structure is generated in .cif format. This structure can be conveniently visualized using the pyvista library, with accompanying data for chromosome and (sub)compartment coloring readily available. Furthermore, distinct .cif structures for each chromosome are extracted, facilitating focused examination of individual chromosomal configurations.

## 3. Discussion

MultiMM offers a user-friendly environment for the 3D modelling of the genome-wide structure of chromatin. Furthermore, the force-field of MultiMM illustrates the existing biophysical knowledge about the folding of chromatin. MultiMM gives an accurate representation for all scales of chromatin, and it allows users to simulate fast and easily the whole chromatin and test different hypotheses about the parameters of the forcefield. Furthermore, MultiMM, offers many functions for fast vizualizations of the structures. The computational time is optimized by running the energy minimization with a graphic card or multiple cores of CPU.

