## Supplemental Materials 1 for "Multiscale Molecular Modelling of Chromatin with MultiMM: From Nucleosomes to the Whole Genome"

In this file we specify the MultiMM model. Initially, we discuss the appropriate datasets that can be used for modelling with MultiMM and the data preparation that is performed within the model. Next, we present the forcefield of MultiMM software, which is separated in four kinds of forces: trivial forces that connect adjacent beads, add stiffness and they perform excluded volume effect, bonded loop forces that model long range loops, compartment or subcompartment forces which represent attracting blocks, and interactions of the polymer structure with lamina. Finally, we present the structure of the software and how it can run.

MultiMM is a multi-scale simulation framework that necessitates a precise definition of scales. This task is particularly challenging due to the inherent variability in most chromatin scales, despite the well-defined lengths of certain chromatin features, such as the radius of nucleosomes. Additionally, simulations are conducted on computers that employ dimensionless units, further complicating the process. OpenMM [1] aids in the definition of these units, but the change in granularity within the simulation results in altered scales. Consequently, 1 nm in OpenMM may not correspond directly to 1 nm in a biological system. To mitigate this issue, we will utilize simulation units (su) instead of nanometers and strive to maintain consistency in the ratios across different chromatin scales at least in order of magnitude approximation. We will refer to nanometers only when we speak about the dimensions of the real biological systems.

### 1. DATA PREPARATION

Firstly, MultiMM imports loop data. The only restriction for these data, is to be in `.bedpe` format. The first three columns should determine the chromosome and genomic coordinates of the left anchor, whereas the next three the coordinates of the right anchor. The seventh column is the strength of the loop $S_{i,j}$, where $i$ and $j$ refer to the anchor locations. Having imported the loops, MultiMM converts them in genomic coordinates, and aggregates the strength of loops with identical indexes by imputing with the average. Then there are two options of modelling, the simplest one is to model all loops with fixed equillibrium lengths $d_{i,j} = 0.1\ su$ or alternatively to correlate the equillibrium length as inversely proportional to the loop count $d_{i,j} \sim 1/S_{i,j}^{2/3}$. The equillibrium lengths of the loops are normalized to be within the interval of $0.1\ su < d_{i,j} < 0.2\ su$. Loop data are essential to run the software, without them it returns error.

Apart from looping data, MultiMM is capable of importing compartmentalization data as well. The user needs to provide a `bed` file with the coordinates of all regions and their (sub)compartment label. The `bed` file is needed to have four columns with the chromosome of interest, the start, end of the region and the compartment label in the third column (A and B). Alternatively, the user can provide subcompartments as input as well (A1, A2, B1, B2). Having imported (sub)compartments, MultiMM generates an array with epigenetic spins $s$ which takes values $s \in \{-1, 0, 1\}$ in case of compartments and $s \in \{-2, -1, 0, 1, 2\}$ in case of subcompartments, with the positive values to correspond to A compartments. A useful software that can be used for the generation of subcompoartemnts from Hi-C data is Calder [2].

### 2. INITIALIZATION OF THE SIMULATION

To set up the simulation, the user needs to provide a configuration file. Within the configuration file, the user needs to define the datasets that he uses, and the forces that are enabled. Not all forces are always needed. MultiMM can either model specific regions of the data, or the whole genome. In case, that the user wants to model the whole genome, needs to enable additional forces that are related to interactions with lamina, compartments and chromosomes. In table S1

| Forcefield Name | Default | Specific Region | Genome-wide |
|---|---|---|---|
| Backbone harmonic bonds | Enabled | Yes | Yes |
| Harmonic angle forces | Enabled | Yes | Yes |
| Loop forces | Enabled | Yes | Yes |
| Excluded volume (LJ) | Enabled | Yes | Yes |
| Container force | Disabled | No | Yes |
| Chromosomal blocks | Disabled | No | Optional |
| Compartment blocks | Disabled | Optional | Optional |
| Subcompartment blocks | Disabled | Optional | Optional |
| Ineteraction with lamina | Disabled | No | Optional |
| Attraction of nucleolus | Disabled | No | Yes |
| Nucleosome interpolation | Disabled | Optional | Optional |

**Table S1.** In this table are the names of all forces in simulations. In "Default" column we show if the particular forcefiled is enabled by default. Furthermore, in the other two columns we add a suggestion if the user needs to enable the particular forcefield in case of modelling a specific region or the whole genome.

we can see a summary of the forces that are enabled by default and the forces that are needed in genomewide simulation. In this manner, forces that are connected to the spherical container are essential to reproduce the final structure. On the other hand, there are some optional forces that can be enabled only if the user provides appropriate data. For instance, block-copolymer forces cannot be modelled without data that provide information about the compartments, or nucleosome interpolation cannot run without data that provide information about the nucleosome density (i.e. ATAC-Seq or MNase-Seq).

The default parameters of the forcefield are provided in table S2. These parameters are tested to work well with the genome-wide simulation. We encourage user to experiment with parameters so as to receive results that they agree to their biophysical intuition. In this supplamentary, we provide the definitions of the forces, which may help users to adjust the parameters according to their expectations.

Simulation was tested on Linux Debian, Ubuntu and Red-Hat based operating systems, with CPU, CUDA and OpenCL parallelization. We suggest users to run it in Linux environments. It is possible to run MultiMM in Windows and Macintosh as well, however the users should acknowledge that they may encounter issues. For example, libraries like pybigwig that is used for loading data for nucleosome interpolation, do not work with Windows. Similarly, despite the fact that it is possible to run MultiMM in Macintosh computers, their technology lack of CUDA parallelization, and it may result to longer computational times.

### 3. FORCEFIELD

To present the forcefield of the simulation we must take into consideration that it can be factored in four basic terms,

$$E = E_{\text{pol}} + E_{\text{lamina}} + \sum_{i,j \in \text{loops}} k_{i,j}(r_{i,j} - d)^2 + \sum_{c \in \{A,B\}} E_c \exp\left(-\frac{r^2}{2r_0^2}\right) \tag{S1}$$

where $E_{\text{pol}}$ represents terms that are responsible for the formation of the main structure of the polymer, $E_{\text{lamin}}$ is the interactions with lamina, the third term represents loop interactions, and the fourth one block-copolymer interactions [3]. In this forcefield the are many different parameters. In the following text, we describe the default values of them, however, the user is able to easily change them by configuring the input configuration file.

### A. Polymer interactions

The polymer interactions term includes basic interactions that are responsible for the formation of the polymer structure. Therefore, it can be described by the following terms,

$$E_{\text{pol}} = \sum_i k_b (r_{i,i+1} - \ell)^2 + \sum_i k_s (\theta_{i,i+1} - \vartheta)^2 + \epsilon \left(\frac{\sigma}{r}\right)^{\alpha} \tag{S2}$$

the first term fixes the the distances between adjacent beads. We assume that the default value of the equilibrium distance is $\ell = 0.1$ su, whereas the spring strength is $k_b = 3 \times 10^5$ kJ/(mol · su$^2$). The second term is an angle bond force responsible for the polymer stiffeness. Therefore, we assume that the equillibrium angle is $\vartheta = \pi$ and therefore tends to make the polymer linear. The strength of the angle force is soft $k_s = 100$ kJ/mol. The third term represents a repelling Leonard Jones potential. Usually, we assume that $\epsilon = 100$ kJ/mol and $\alpha = 3$.

| Parameter Name | Symbol | Value | Units |
|---|---|---|---|
| Harmonic Bond Strength | $k_b$ | $3 \times 10^5$ | $kJ/(mol \cdot su^2)$ |
| Harmonic Bond Equilibrium Distance | $\ell$ | 0.1 | $su$ |
| Harmonic Loop Strength | $k_l$ | $3 \times 10^4$ | $kJ/(mol \cdot su^2)$ |
| Harmonic Loop Equilibrium Distance | $d$ | $[0.1, 0.2]$ | $su$ |
| Harmonic Angle Strength | $k_s$ | $10^2$ | $kJ/(mol)$ |
| Harmonic Equilibrium Angle | $\vartheta$ | $\pi$ | radians |
| LJ Strength | $\epsilon$ | $10^2$ | $kJ/mol$ |
| Length Scale of LJ | $\sigma$ | 0.05 | $su$ |
| LJ power | $\alpha$ | 3 | – |
| Container Strength | $C$ | $10^3$ | $kJ/(mol \cdot su^2)$ |
| Lamina Attraction Strength | $B$ | $10^2$ | $kJ/mol$ |
| Attraction of the Nucleolus | $G$ | $10^2$ | $kJ/mol$ |
| Strength of Chromosomal Attraction | $E_n$ | $10^{-3}$ | $kJ/mol$ |
| Chromosomal Scale factor | $k_C$ | 0.3 | $su^{-4}$ |
| Compartment Energy Levels | $E_c$ | $[1, 2]$ | $kJ/mol$ |
| Maximum nucleosomes per bead | $n_{max}$ | 4 | – |
| Nucleosome radius | $r_n$ | $10^{-2}$ | $su$ |
| Points per nucleosome | $p_n$ | 20 | – |
| Zigzag angle | $\phi_{norm}$ | $\pi/5$ | radians |

**Table S2.** Table of default simulation parameters. Values inside brackets represent intervals of values.

### B. Lamina interactions

In general case, we assume that the whole genome is consisted by separate polymer chains that represent chromosomes. The whole chromatin structure is enclosed within a spherical container with radius $R_2$ that represents the wall of nucleus. Furthermore, we assume that there is an empty space with radius $R_1$ in the center of the polymer structure that represents nucleolus. In this perspective, we have two walls and polymer structure lives between them.

Apart from the spherical boundary walls, we have another two types of interactions with lamina: attraction of B compartment to lamina, and attraction of smaller chromosomes to the nucleolus. Consequently, this potential has the following form,

$$E_{\text{lamina}} = E_{\text{bc}} + E_{\text{Bl}} + E_G \tag{S3}$$

where the first term represents the lamina boundary condition in the walls $R_1$ and $R_2$,

$$E_{\text{bc}} = C\left(\max(0, r - R_2)^2 + \max(0, R_1 - r)^2\right) \tag{S4}$$

where $C$ is the strength of the wall and it is given the value $C = 1000$ kJ/(mol · su$^2$) by default. The radii $R_1$ and $R_2$ can be correlated with the simulation length $L$ so as to be $R_1 = 0.5 \times (L/(5 \times 10^4))^{1/3}$ $[su]$ and $R_2 = 8R_1$. Therefore, $R_2$ represent the outer boundary of the nucleus lamina, whereas $R_1$ represents the inner nucleolus [4, 5]

The second term represents the attraction of B compartment with lamina [6–8]. This term becomes minimum near the walls $R_1$ and $R_2$, and thus it has the following functional form,

$$E_{Bl} = B\left(\sin^8\left(\frac{r - R_1}{R_2 - R_1}\right) - 1\right)(\delta(s+1) + \delta(s+2)) \tag{S5}$$

where $\delta(\cdot)$ represent delta Knonecker function which becomes 1 only when the compartment index $s$ is negative and therefore the region is within the B compartment. On the right-hand-side of the equation there are $\Theta(\cdot)$ Heaviside step functions which make the potential zero in the region that it is not within the container. We set $B$ to be by default 100 kJ/mol representing a soft boundary effect.

Finally, we can model attraction of smaller chromosomes [9–11] with nucleolus which is represented by the wall $r = R_1$, by writing down the following potential,

$$E_G = Gs_c\left(\sin\left(\frac{r - R_1}{\ell_G}\right) - \left(\frac{r - R_1}{\ell_G}\right)^2\right) \tag{S6}$$

where $s_c$ takes values in the interval $[0, 1]$ and it is 1 for the chromosome with smaller length and 0 for the chromosome with the larger length, and $G$ takes small values like 100 kJ/mol. By making $G$ stronger, it can result to very intense attraction to nucleolus, and thus we suggest to the users to keep the value of this parameter low.

#### C. Looping interactions

We described the loop potential as following,

$$E_{\text{loop}} = \sum_{i,j \in \text{loops}} k_{i,j}(r_{i,j} - d)^2 \tag{S7}$$

where $i$ and $j$ are locations where we know from the experiment that they are in close proximity due to some loop force. It is needed to have at least loops from experiment so as to run MultiMM. The loops should be in the format of a `.bedpe` file. The equillibrium length $d_{i,j}$ as inversely proportional to the loop count $d_{i,j} \sim 1/S_{i,j}^{2/3}$. The equillibrium lengths of the loops are normalized to be within the interval of 0.1 $su < d_{i,j} < 3 \times 0.2$ $su$. The harmonic bond strength $k_l$ is always fixed $3 \times 10^4$ $kJ/(mol \cdot su^2)$.

#### D. Block-copolymer interactions

We described compartment forces with the block copolymer potential [3]

$$E_{\text{block}} = \sum_{c \in \{A,B\}} E_c \exp\left(-\frac{r^2}{2r_0^2}\right) \tag{S8}$$

where we assume that $r_0$ is the proximity of the long range compartmentalization interaction, and $E_c$ are the energy states for compartments $c$. Depending on the data that we have, we may have compartments (i.e. $A$, $B$) or subcompartments (i.e. $A1$, $A2$, $B1$, $B2$). In case that we have compartments and $s \in \{-1, +1\}$ then we can write

$$E_c = E_a\delta(s_1 - 1)\delta(s_2 - 1) + E_b\delta(s_1 + 1)\delta(s_2 + 1) \tag{S9}$$

whereas in case that we have subcompartments $s \in \{-2, -1, 1, 2\}$ then we can generalize energy states to four,

$$\begin{aligned} E_c = &E_{a1}\delta(s_1 - 2)\delta(s_2 - 2) + E_{a2}\delta(s_1 - 1)\delta(s_2 - 1) \\ &+ E_{b1}\delta(s_1 + 1)\delta(s_2 + 1) + E_{b2}\delta(s_1 + 2)\delta(s_2 + 2), \end{aligned} \tag{S10}$$

where the energy levels are set so as to have stronger attraction for B compartments which are linked to more condensed regions. Therefore, we can set the parameters $E_a = -1$ kJ/mol and $E_b = -2$ kJ/mol for compartments or $E_{a1} = -1$ kJ/mol, $E_{a2} = -1.33$ kJ/mol, $E_{b1} = -1.66$ kJ/mol and $E_{b2} = -2$ kJ/mol in case that we have subcompartments. The default proximity parameter for compartmentalization is $r_0 = (R_2 - R_1)/20$.

In case that there are not enough loop interactions (eq. S7) in our dataset and most of the loops have small lengths or compartmentalization data are missing, then we can use a force to make chromosomes more globular. This force does not have a direct biological meaning and this is why it is disabled by default. Therefore, there is another one block-copolymer kind of potential that operates over chromosomes and it is disabled by default. This potential can be described by the equation,

$$E_{\text{chrom}} = \sum E_n(k_C r^4 - br^3 + cr^2) \tag{S11}$$

where $k_C = 0.3\ su^{-4}$ is a parameter that controls how flat is the potential near the minimum, and $E_n = 10^{-3}$ kJ/mol. By controlling the parameter $E_n$ we can have very loose chromosome when it is close to 0 and very globular when it is close to 1. We suggest modeling with small values of the energy, since chromosomes are considered loose. Parameters $b = 1\ su^{-3}$ and $c = 1\ su^{-2}$ exist in equations only for the complete definition of units.

| Parameter Name | Symbol | Function |
|---|---|---|
| Small Container Radius | $R_1$ | $R_1 = 0.5 \times (L/(5 \times 10^4))^{1/3}$ |
| Big Container Radius | $R_2$ | $R_2 = 8R_1$ |
| Std of Block-Copolymer Interactions | $r_0$ | $r_0 = (R_2 - R_1)/20$ |
| Equillibrium Loop Length | $d_{i,j}$ | $d_{ij} \sim 1/S_{i,j}^{2/3}$ |

**Table S3.** Table of simulation parameters that do not take fixed values and are fucntions of other parameters.

### E. Nucleosome Interpolation

Nucleosome scale is included as an interpolation method in MultiMM by assuming a beads-on-the-string model of nucleosomes [12, 13]. Therefore, it is assumed that DNA wraps around histones 1.6 times like a counter-clockwise helix. The user imports the ATAC-Seq signal, and the simulation places nucleosomes in a coarse-grained manner [14, 15]. Therefore, regions with weak ATAC-Seq signal will correspond to nucleosome rich regions, whereas regions with high ATAC-Seq signal would be nucleosome poor. For the purpose of this study, we used p-value of ATAC-Seq data which follows an inverse relation with the density of nucleosomes, because low p-value means that there is a significant difference in chromatin accessibility, suggesting changes in nucleosome density or other regulatory mechanisms. Therefore, regions with high ATAC-Seq p-value correspond to nucleosome dense regions, and low ATAC-Seq signal. Consequently, we aggregate the signal so as to fit the simulation dimension by computing the average ATAC-Seq p-value for each simulation bead. The logarithm of the signal is then normalized to $[0, 1]$ and used as a basis for the number of nucleosomes to align in a given chromatin segment. The number of nucleosomes is calculated as proportional to that normalized signal with the maximum number assigned to the segments with the strongest signal and no nucleosomes are located in the segments with the weakest. The user can specify the maximum number of nucleosomes that are allowed to be in a single bead.

In order to initialize the nucleosome positions the generic nucleosome helix was first created. This generic helix was positioned in the particular DNA fragments specified by two points in 3D space: the start $p_1$ and end $p_2$ of the nucleosome helix. The function translates the generic helix coordinates defined in standard basis of a coordinate vector space into the basis defined by a vector between start and end point of the specified nucleosome position ($\vec{p_1 p_2}$) and two other orthogonal vectors. The second vector from this base is defined by a perpendicular component of the vector ($\vec{p_0 p_1}$) where $p_0$ is a point lying on an original segment corresponding to a given nucleosome. Third vector is found by cross product of the other two and all of them are normalized to form a new orthogonal basis. The coordinates of the original generic helix are then written down in the coordinates of this basis with the starting point of $p_1$, producing a desired helix positioning along the DNA strand. This positioning is defined by a beads-on-a-string or zigzag model with linker DNA between nucleosomes set to 3.45 times the nucleosome radius. Firstly, on each segment equidistant points are determined, one point per nucleosome. From each point the nucleosome is going to be located at half the distance of the linker DNA away from the segment by the function described above. Consequent nucleosomes are located on the opposite sides of the segment with the additional small random angle thus creating a structure roughly corresponding to the zigzag model. The angle of the last nucleosome in a segment is passed to the next assuring that the consequent nucleosomes on a different segment will not be placed on the same side of the segment.

The nucleosome helices are initialized based on a couple of assumptions about the nucleosome dimensions. Firstly, it is assumed in our simulation that chromatin wraps around nucleosome 1.65 times [16]. Secondly, the ratio between the nucleosome radius and its height is assumed to be 1 based on the estimates in [17] that the nucleosome structure has a height of about 5.5 nm and a diameter of 11 nm. The ratio between the nucleosome radius and average linker DNA length is set to 1/3.45, which was based on the estimate of the average linker DNA length of 55 bp [18] and the length of a single DNA base pair of 0.34 nm. Finally, the angle $\phi_{norm}$, for which the angle through which every second nucleosome is rotated in a zigzag configuration, is by default set to $\pi/5$ [19].

Furthermore, it is worth comparing the length of the nucleus radius with the scale of loops. The sizes of loops can be highly variable, from 50 *kb* to 200 *kb* [20], where each *kb* can be considered as a 0.34 *nm* DNA fiber. This means that the average ratio of loop length over nucleosome radius is on the order of magnitude of $10^2 - 10^3$, because in simulation we exclude too small loops that cannot be modeled with long-range forces. In our simulation, we assume that a single loop consists of tens of beads of 0.1 *su*, and the radius of a nucleosome is $10^{-2}$ *su*. Considering that each loop consists of tens or hundreds of beads, the ratio remains similar to the real one.

### 4. OPTIMIZATION

For the optimization of the model there are many different options of OpenMM integrators that are included in the model: Verlet and Langevin integrator with fixed and variable steps, browian integrator, and AMD integrator. In principle, it is enough to run an energy minimization to produce a reasonable structure. In case that the users are not user-specified satisfied with that, they implement relaxation dynamics, by letting the system to evolve for some simulation steps. The software outputs the energy and temperature of the simulation and the user and diagnose if the simulation works correctly.

### 5. RUNNING MULTIMM

MultiMM has two modelling modes: modelling the whole genome or modelling a specific region. In the first case, the user does not need to specify any region, but needs to provide data for all chromosomes of the simulation. In the second case, the user needs to specify the modelling region including the chromosome index and the coordinates in units of genomic distance. All the parameters and input data are provided in a configuration input file. In GitHub repository there is a complete manual of the parameters of the model and examples about its usage. Apart from data and parameters, the user should also specify the initial structure. For genome-wide simulations, we propose Hilbert curve as an initial structure because the simulation needs less time to minimize the energy since the structure is already folded.
